# Choosing between personal values: The Pavlovian substrates of intrinsic preferences

**DOI:** 10.1101/856294

**Authors:** Roberto Viviani, Lisa Dommes, Julia Bosch, Petra Beschoner, Julia C. Stingl, Tatjana Schnell

## Abstract

Several brain circuits interact in computing the value of choices between options, as when we express our preference between a set of available consumer goods. Here, we used a procedure developed in functional neuroimaging studies of consumer choice to identify the neural substrates activated by choosing between values that, when put into practice, can give meaning to one’s life, such as achievement, community, tradition, or religion, and are unrelated to material needs or financial security. In a first sample (N=18), instead of the neural substrates usually associated with choice between consumer goods, we found activation of the amygdala, a limbic system structure which presides over assignment of values to stimuli according to immediate affective experience and promotes responses according to their association with potential rewards. This unexpected finding was replicated in a second independent sample (N=18). These results are consistent with views arguing for the existential nature of values that give meaning to one’s life here and now, in contrast to maximizing long-term utility.

## Introduction

A unifying notion emerging in the behavioural study of choice is that of ‘value’ (Glimcher and Fehr 2014). Like the economic notion of ‘utility’, value underlies choices by integrating all relevant information about the alternatives under consideration into a single quantity on a subjective preference scale. Such a preference scale is required in a rational agent, since making consistent choices is logically equivalent to maximizing a single utility function that orders options according to their desirability (Samuelson 1938). Studies in laboratory animals and in humans have shown that a neural signal with many properties of an integrated subjective value signal is present in a brain network system centred on the medial orbitofrontal cortex (Montague and Berns 2002; Padoa-Schioppa and Assad 2006; Levy and Glimcher 2012). This system, known as *goal-directed* (Dickinson and Balleine 2002), is active in humans when choosing between consumer goods of very different types (Rangel and Hare 2010; Bartra et al. 2013; Clithero and Rangel 2014), including food and money (FitzGerald et al. 2009; Smith et al. 2010; Levy and Glimcher 2012).

In both animals and humans, however, there seem to be multiple competing systems that compute values (Daw and O’Doherty 2014), a finding that may contribute to explaining why human behaviour sometimes only approximates the consistent choices of a rational agent (De Martino et al. 2006; Weber et al. 2007; Balleine et al. 2008). This view is based on empirical evidence on the involvement of other brain structures in the computation of subjective values. One such alternative brain structure is the amygdala (Murray 2007; Seymour and Dolan 2008; Morrison and Salzman 2010; Janak and Tye 2015), which is involved in the forming of associations of conditioned stimuli with the properties of the original rewards, including their motivational and affective valence (Everitt et al. 2003; Balleine and Killcross 2006). Because of this role, values computed in the amygdala are often referred to as *Pavlovian values* (Dickinson and Balleine 2002; Daw and O’Doherty 2014). Pavlovian values appear to arise in direct connection with affective experience (Murray 2007; Seymour and Dolan 2008; Morrison and Salzman 2010). In contrast, and in keeping with their utility-like character, goal-directed values observed in the orbitofrontal cortex not only reflect past memories of rewards, but can also give rise to choices that can take into account the expected future outcomes of decisions, thus integrating different factors in computing preferences (Bechara et al. 1999; Hampton et al. 2006; Noonan et al. 2010; McDannald et al. 2012; Rudebeck and Murray 2014; Wilson et al. 2014).

Little is known about the mechanisms through which preferences are formed among human ‘values’ as this term is used in common parlance, i.e. properties of objects or actions that can give a sense of purpose and meaning to one’s life. In the present study, we used the procedure based on eliciting a gradient of preference in neuroeconomic studies of binary consumer choice to ask participants in the scanner to choose between situations according to the importance they had in their life (Figure 1, left). The available alternatives (two at a time) were drawn from all combinations of a set of six representative values that empirical research has identified as sources of meaning in life, when put into action (community, nature, knowledge, religion/spirituality, achievement, tradition; Schnell 2011). These values may be understood as intrinsic, as worthwhile per se and not in terms of possible benefits arising therefrom. To have a term of comparison (‘functional localizer’), we also asked participants to choose between snacks in a separate run in the same scanning session (Figure 1, right). This task has been shown in the neuroeconomic literature (Rangel and Hare 2010; Bartra et al. 2013; Clithero and Rangel 2014) to reliably activate the medial orbitofrontal cortex, a key substrate for the representation of goal-directed values that has been associated with the computation of economic utility (Padoa-Schioppa 2011; Levy and Glimcher 2012).

**Figure 1.**
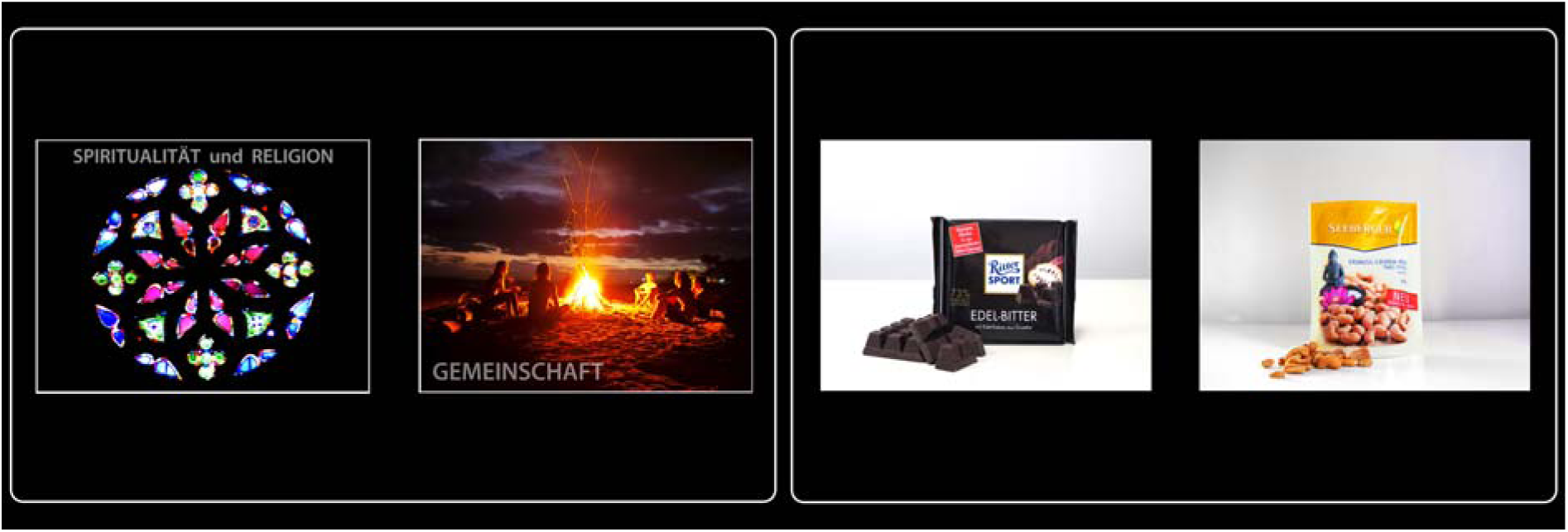
Examples of trials where participants chose between personal values (left) or snacks (right). Personal values were presented two at a time in all combinations available from the set of six values. The figures referring to personal values contained no specific religious symbols, possible allusions to substance of abuse, definable human bodies or parts thereof that might have been sexually arousing, or identifiable faces or facial expressions.

One way of looking at personal values is that they refer to long-term goals followed during a whole or large part of a lifetime (Emmons 2003; Baumeister et al. 2013), suggesting a possible role of goal-directed processes, as these latter support decisions based on future consequences. Furthermore, a virtuous life seems to require the capacity to sustain long-term goals (Thaler and Shefrin 1981; Metcalfe and Mischel 1999), in contrast to the short-term satisfaction based on hedonic pursuits (Ryan and Deci 2001; Huta and Ryan 2010). However, personal values can also imbue life with meaning on a daily basis (Cantor et al. 1991), an aspect of human experience that is at present poorly understood (Schnell 2020). This issue is also of importance to understand pathological states in which motivational priorities are affected, as when a patient suffering from depression reports that she can no longer find comfort in religion, even when it had previously provided direction to her life.

Functional neuroimaging studies in man have shown the involvement of areas belonging to the goal-directed orbitofrontal control circuit when exercising self-control in opting for a larger but more distant reward (Kable and Glimcher 2007). Furthermore, self-restraint in goal-directed control may also recruit circuits associated with effortful attentional processes or executive function (McClure et al. 2004; Cohen 2005; Hare et al. 2009), which may cooperate with value circuits to achieve self-regulation. A primary issue of interest of the present study was therefore to see if we could find evidence of recruitment of goal-directed control in the prefrontal cortex when participants were asked to express preference among personal values.

## Results

In their responses during the experiment, all but two participants expressed preferences for the values nature and community, which achieved top preference scores, while religion/spirituality and tradition were regarded with disfavour by most participants. This pattern corresponds to previous findings in young German-speaking individuals of this age and social extraction (Schnell 2016; Schnell and Becker 2007), indicating that participants were basing their decisions on preferences between the personal values depicted in the trials. Because all options were systematically paired, the maximal preference score obtained in each participant is related to the consistency of choices. Mean max scores were slightly higher in the personal values than in the snacks sample (5.2 and 5.0, respectively), possibly indicating stronger preferences in the personal values data. This difference, however, was not significant (Wilcoxon signed rank test for paired samples, p = 0.18).

To elicit the signal of comparative valuations, we regressed the brain signal of choice trials on the gradient of preference, a quantity that appears as the key variable in formal models of optimal information sampling in binary choice (Bogacz et al. 2006; Krajbich et al. 2010) and was found in neuroimaging studies to reflect valuations in the medial orbitofrontal cortex (Serences 2008; Boorman et al. 2009; FitzGerald et al. 2009; Hunt et al. 2012). The score of how many times each personal value was chosen was computed in each participant, and the difference between the scores of the two presented alternatives calculated for each trial. The interaction between these difference scores and a regressor modelling the activity of trials relative to baseline was added to the model (‘parametric modulation’), therefore capturing variance associated with the gradient of preference and not due to task effects common to all trials.

Contrary to our hypothesis, no involvement of the medial orbitofrontal cortex was observed in association with the comparative valuation of personal values, even if this area (as defined in Levy and Glimcher 2012) was active as expected when regressing the data on the gradient of preference in the snack data (Montreal Neurological Institute coordinates −14, 34, −10, BA11, *t* = 3.84, *p* = 0.001; Figure 2, bottom row). This null finding persisted also when using the effect of snack choice to localize the voxels for the signal associated with comparative valuation. Instead, statistical analysis revealed the bilateral involvement of the amygdala, especially in its basolateral portion (20, −4, −22, *t* = 3.95, *p* = 0.001; −26, −2, −26, *t* = 3.87, *p* = 0.001, both uncorrected; Figure 2, top row). In the amygdala there was no signal associated with the comparative evaluation of snacks, even at uncorrected significance levels. No activation in the amygdala was observed in association with the sum of the scores of the displayed alternative personal values (instead of the difference).

**Figure 2.**
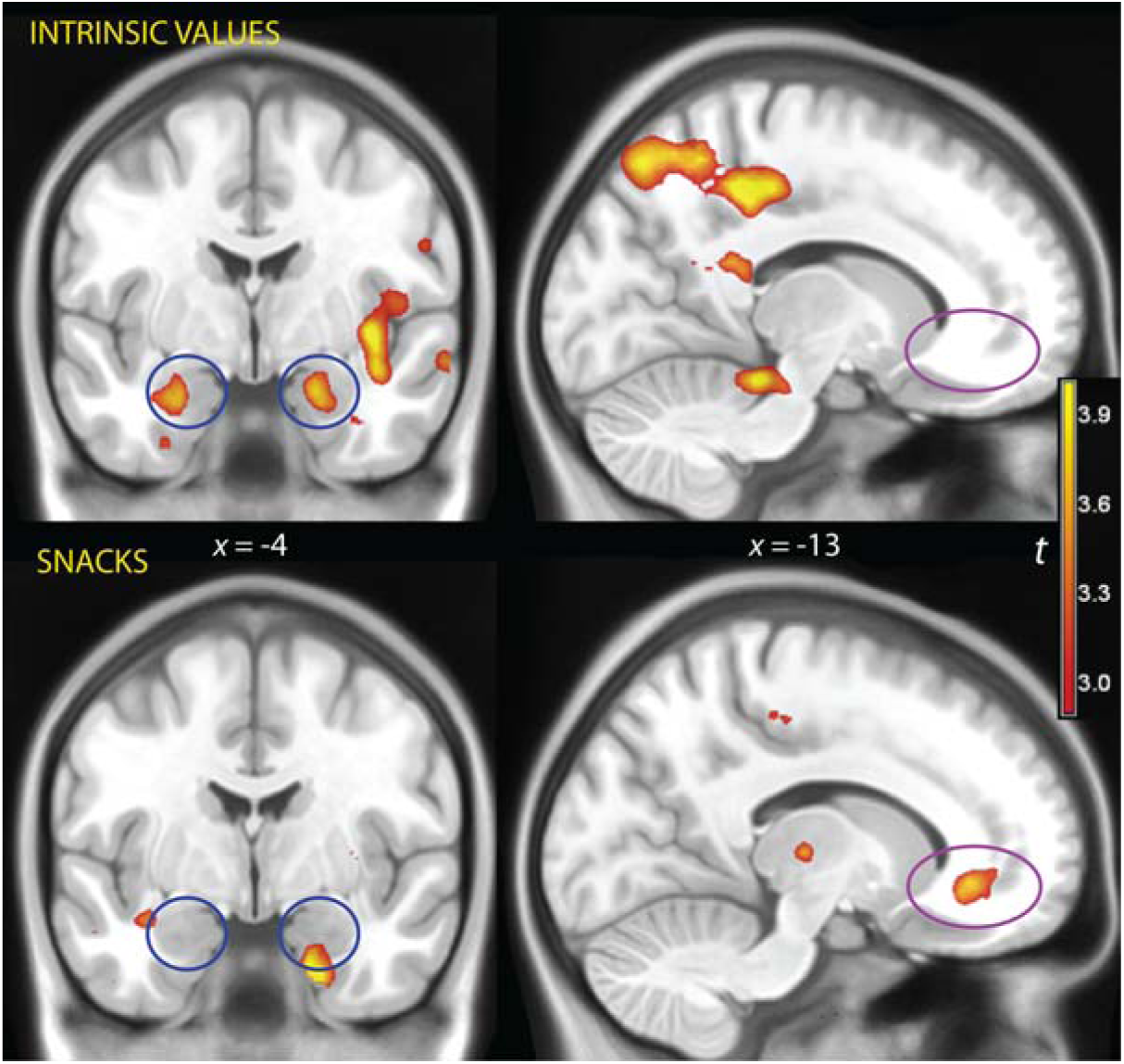
Top row: when choosing between personal values, participants activated the amygdala bilaterally when the gradient of preference was large (blue circles). No such activation was detected when choosing between snacks (bottom row). In contrast, choice between snacks activated the orbitofrontal cortex (magenta circle) in the region described in the literature to be associated with the computation of a comparison value signal (Levy and Glimcher 2012). Data represent *t* values thresholded for illustration purposes at p < 0.005, uncorrected, overlaid on a template brain. MNI coordinates in mm.

Other areas that were significantly activated by the comparative valuation of personal values were the supramarginal gyrus in the inferior parietal lobe on both sides (−60, −30, 42, BA40/2, *t* = 6.59; *k* = 2091, *p* = 0.018; and 64, −24, 46, BA40/2 *t* = 3.89, *k* = 3097, *p* = 0.015, both cluster-level corrected). These bilateral clusters of activation extended medially into the middle cingular cortex (18, −38, 44, *t* = 4.67; −16, −38, 42, *t* = 4.89, Figure 2, top right) and ventrally into the temporoparietal junction (−56, −32, 28, BA40/42, *t* = 3.85; 50, −36, 38, BA40, *t* = 3.79) and the left superior temporal gyrus (−66, −30, 20, BA22, *t* = 3.64). On the right, this latter activation extended into the superior temporal gyrus and the insula (40, −8, −12, BA20, *t* = 4.83) but, as a separate cluster, failed to reach significance after correction. No effect of the gradient of preference in the ventral striatum was observed, even at uncorrected levels.

Because the activation in the amygdala was an unexpected finding that failed to reach significance after correction, we conducted a replication study of the values decision task where we designated the amygdala as a region of interest a priori. In this second study, we replaced the snacks decision task with one where participants decided about pet food alternatives to check for the activation of the amygdala related to possibly aversive items. In the value decision task, this second study replicated the finding of a signal correlated with a gradient of preference in the amygdala (−26, −4, −26, *t* = 3.48, *k* = 23, *p* = 0.067, and 24, 0, −24, *t* = 3.38, *k* = 52, p = 0.031, cluster-level corrected for the region of interest), and the failure to elicit a corresponding signal in the orbitofrontal region and in the ventral striatum. The null findings in the medial orbitofrontal cortex did not change when we pooled data from both studies. In contrast, in the pooled data the effect in the left amygdala reached peak-level significance when corrected for the whole brain (−26, −4, −24, *t* = 5.03, *p* = 0.034). A similar improvement in the evidence for an effect was observed in the right amygdala (24, −2, −26, *t* = 4.73, *p* = 0.076, same correction).

To characterize the course of the signal independently from the BOLD modelling strategy of the trial, we fitted a Fourier series to the data at the first level adjusted only for the effects of the intercept and the movement covariates (Figure 3). The course of the signal in the low and high evidence trials shows that the effect of the gradient of preference ensued gradually during the trials and peaked after the decision. Although the signal rose during all trials, the rise was steeper in the high gradient trials.

**Figure 3.**
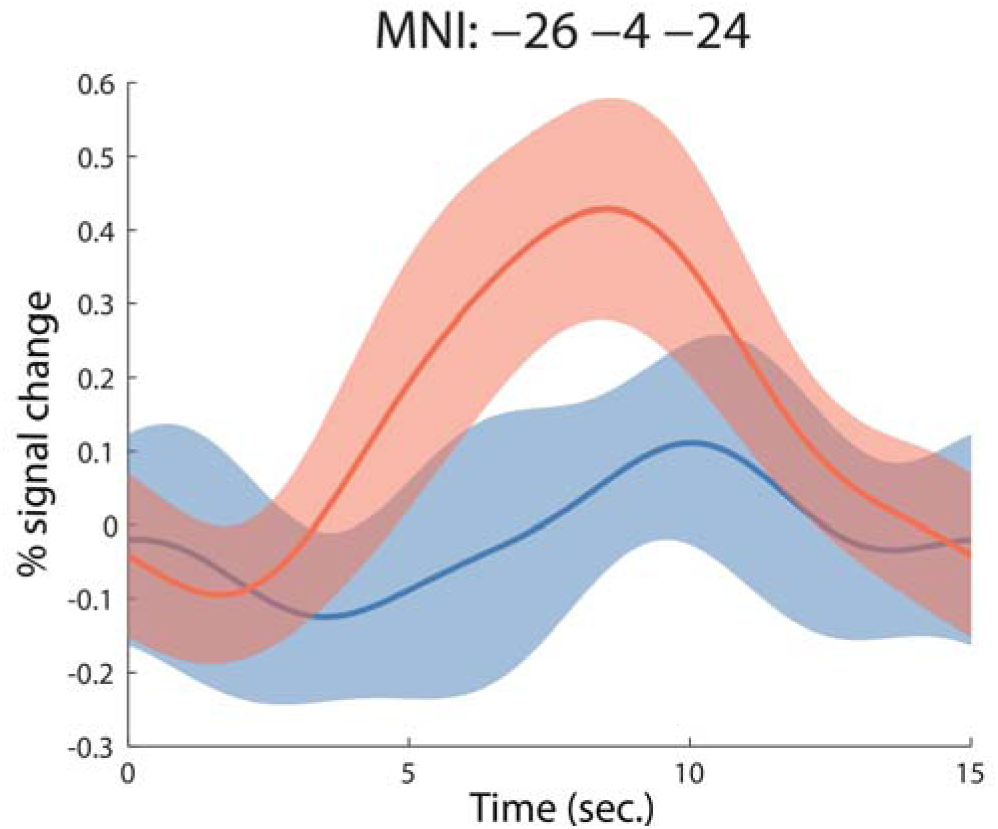
Plot of the signal course of trials in the highest (red) and lowest (blue) tertiles of the gradient of preference in the left amygdala (95% confidence intervals). Time 0 corresponds to the beginning of the trial, when the alternative options were presented. The BOLD response is expected to rise with a 2-3 sec. delay relative to the timing of the stimuli.

To investigate the possibility of an effect of choices to which participants may have felt no personal relation or might be perceived as aversive, we considered only the trials in which no item had received a preference score of one or less (given that the possible scores ranged from zero to five, this cut-off excluded trials containing the two least preferred items from the set of six. This appeared to be a reasonable cut-off given that the values were generally positively connotated). The aim was to create a cleansed sample where choices were made on the basis of appetitiveness or neutrality, not aversion. Because there were a smaller number of such trials and the gradient of preference was smaller, we used the pooled data from both studies. Even in these cleansed data, the gradient of preference was associated with an activation of the amygdala (18, −6, −24, *t* = 3.42; −28, −8, −26, *t* = 2.41), suggesting that the presence of aversive or personally irrelevant values was not responsible for the reported effect.

Information on choices computed on the basis of aversion, to the judgment that some choice may be inappropriate, or to choices taken on alternatives to which participants could not relate personally, was also provided by the pet food choice task of the second study. While pet food items may not be entirely unfamiliar, they are likely to be mildly aversive. It may also have been difficult for participants to form self-attributions of preferences between them, as between values toward which they may have felt indifferent. However, there was no activation of the amygdala associated with the gradient of preference (least aversion) between pet food items (no voxel over uncorrected significance levels).

## Discussion

In laboratory animals, the amygdala appears to play a key role in learning and representing values of immediate affective or motivational significance (Murray 2007; Morrison and Salzman 2010), be they aversive or appetitive (Paton et al. 2006). These findings underlie views that attribute a key role to the amygdala in preference and choice (Murray 2007; Morrison and Salzman 2010). However, in neuroimaging studies of choice in humans, activation of the amygdala is uncommon, in striking contrast with the robust recruitment of the medial orbitofrontal cortex (Bartra et al. 2013; Clithero and Rangel 2014). This suggests that in adults of our species explicit consideration of the consequences of one’s actions, as represented in goal-directed choice, is the common means to make decisions (Bechara et al. 1999; Hampton et al. 2006; Rudebeck and Murray 2014; Wilson et al. 2014). Amygdala involvement is usually detected by introducing circumstances that perturb choices, such as framing choices with different enticing descriptions (De Martino et al. 2006; Weber et al. 2007; Grabenhorst et al. 2013) or manipulating the current desirability of options (Gottfried et al. 2003). Accordingly, goal-directed processes have been viewed as those that best satisfy the requirements of normative theories of choice to ensure consistency of expressed preferences in humans, while Pavlovian values such as those represented in the amygdala have been prevalently associated with behavioral anomalies or biases (Dayan et al. 2006). In the present experiment, however, the signal in the amygdala was associated with the comparative valuations directly, providing evidence for the long-standing suggestion of a role of this structure in determining choices also in humans (Seymour and Dolan 2008), but only when specific types of preferences are at stake. In contrast, activation of the prefrontal neural substrates of goal-directed and executive processes, which are often associated in the literature with adaptive choice, self-control, and the capacity to sustain long-term objectives (Emmons 2003; Baumeister et al. 2013; Huta and Ryan 2010; Vötter and Schnell 2019), was inconsistent or absent. These data suggest that to make choices among personal values, humans use a network that is often reported in studies of choice in laboratory animals, while to make mundane choices, humans use more refined goal-directed mechanisms.

Neuroimaging studies in man also show the amygdala to be frequently active in circumstances where it may be associated with the emotional salience of stimuli (Whalen et al. 2002; Dolan 2002; Phelps and LeDoux 2005), including potential rewarding choices (Arana et al. 2003). However, an interpretation of the present data in terms of salience (or of any another sensory property of the stimuli) is unlikely, since in the regression of Figure 2 trials containing image pairs of personal values with a high gradient of preference were contrasted with trials presenting the same images, differing only in the way in which the pairs were combined. In confirmation of this reasoning, it has been shown that the amygdala is not active as a correlate of evidence for choice-making even with emotionally arousing stimuli. For example, no activation of the amygdala was detected in association with the gradient of scores when participants were asked to decide which of two displayed faces experienced the most intense emotion (Viviani et al. 2018), even if faces with emotional expressions are potent activators of the amygdala.

The involvement of the amygdala suggests that choice in the domain of personal values may not be best thought of as maximizing value through consideration of the long-term consequences of choices or goal-directed representations of outcomes, as we initially hypothesized. Instead, priorities among personal values might be at least partially maintained by a system representing their affective desirability (or lack thereof) at the time the decision was taken. This finding is consistent with views that emphasize the importance of personal values in imbuing life with meaning on a daily basis (Cantor et al. 1991; Hammell 2004), and define lifetime goals within an existential framework by emphasizing their difference from the maximization of a utilitarian criterion (Schnell 2009, 2020). Furthermore, it is consistent with the conjecture that in humans the amygdala may mediate affective sources of preferences and biases associated with ideas, aspirations, or convictions (Murray 2007). The involvement of affective neural substrates may also explain why personal values figure prominently among the motives of individuals that report high levels of meaningfulness in their lives (Schnell 2011).

## Methods and Materials

The study was conducted at the Psychiatry and Psychotherapy Clinic of the University of Ulm, Germany, and was formally approved by the Ethical Review Board of the University (applications nr. 01/15 and 122/17). Participants without history of psychiatric or neurologic disorders were recruited through local announcements and admitted to the study after providing written informed consent. To characterize the sample, we administered self-rating scales for depressiveness and anhedonia (CES-D, depressiveness: Radloff 1977, German version: Hautzinger and Bailer 1993; SHAPS-D, anhedonia: Snaith et al. 1995, German version: Franz et al. 1998). We also administered the meaningfulness scale from the Sources of Meaning and Meaning in Life Questionnaire (SoMe, Schnell and Becker 2007; Schnell 2009, 2014), a validated scale to measure the degree of subjectively experienced meaning in life, operationalized as a fundamental sense of purpose, orientation, coherence, and belonging. Both study samples were composed of 18 participants (first study: age 24.4±4.8, 4 males; CES-D scores 8.6±6.0, SHAPS-D 21.4±5.0, mean SoMe meaningfulness 3.4±1.0; second study: age 23.2±4.2, 4 males; CES-D scores 5.7±6.0, SHAPS-D 20.8±4.4, mean SoMe meaningfulness 3.7±1.8). In the combined sample where only the trials were considered when the values on display had a score of two or more, some participants were excluded where there were less than four such trials (this happened if a participant gave to 3 items a score of 1). This left 24 participants for the combined sample.

In the task where participants chose between personal values, participants were asked to select the option that corresponded best to what would be felt to be purposeful and to give meaning in their life. In the task where participants chose between snacks (first study), they were informed that they would receive the most preferred snack after the experiment (which they did). In the task where participants chose between samples of pet food (second study), they were told to imagine that they would receive the sample after the experiment and that they would have to eat it.

In each trial, participants were asked to choose among the alternative items displayed on screen (two at a time) by pressing a button on the side of the chosen alternative. Button presses indicating decisions could be made only after 2.5 sec of display. This time was cued by the appearance of two blue circles symbolizing the buttons under the depicted items. At the appearance of the cues, participants had 1.5 sec to make their choice, after which the trial terminated. Trials were separated by variable intervals averaging 10.5 sec determined with a random exponential schedule bounded at 9 and 12 sec. Each trial displayed one of all possible combinations of the six items available for choices (personal values or snacks in the respective sessions), giving 15 trials for a session duration of 3 min 40 secs. The trial images were displayed using standard software (Presentation 14, Neurobehavioral Systems Inc., Albany, CA) through goggles masking the whole field of vision (Resonance Technology Inc, Northridge, CA). The images avoided representation of faces or body parts, but contained short written text (descriptive names of personal values and the labels of the snack products on the packages). Participants were told to choose between personal values according to their relative personal importance, and between snacks according to what they would like to consume. Participants familiarized with the task in a brief session prior to scanning.

In adopting the retarded response design, we intended to avoid large systematic differences in the duration of the trials, since shorter reaction times are known to arise with larger preference gradients (Bogacz et al. 2006). This would have led to a possible confound in the estimated effects of the BOLD signal due to systematic differences in trial duration in association with preference gradients. In the present study, qualitatively identical results were obtained by modeling trials using a fixed duration and using the effective durations of the trials (here, only the fixed duration model was shown for brevity).

Previous functional MRI studies have shown the signal associated with the gradient of choice to arise in the data when fixating the chosen item (Lim et al. 2011). Fixation to the chosen item converges sooner (and hovers longer) when the gradient is larger. This suggests a relationship between the signal and quicker convergence of a drift diffusion sampling process mediated by fixation (Krajbich et al. 2010). If this model is correct, inducing similar trial durations by enforcing choice at the end of the trial samples the accumulated signal arising from fixating the chosen item after convergence to the chosen item, thus reflecting the steepness of the gradient of preference.

Data collection and analysis were identical in the personal values and snacks sessions. MRI data were obtained with a 3-Tesla Magnetom Prisma (Siemens, Erlangen, Germany) MRI system equipped with a 24-channels head coil. Structural images were obtained and screened to exclude pathology. Functional MRI data were acquired with a gradient-echo echo-planar imaging (EPI) sequence (TR/TE 1970/36 msec., flip angle 90°, bandwidth 1776 Hz/pixel, 32 transversal slices, anterior-to-posterior phase encoding, FOV 192/174 mm in the frequency/phase encoding directions, giving an image size of 64×58×32 voxels). Slice thickness was 2.5 mm with a gap of 0.75 mm, giving a voxel size of 3×3×3.25 mm. In each session, 114 images were acquired.

Data were analyzed with a general linear model (Friston et al. 1995) using the software SPM8 (publicly available at http://www.fil.ion.ucl.ac.uk/spm/). After realignment and normalization in MNI coordinate space (default parameter settings, resampled voxel size 2×2×2 mm) and smoothing (isotropic Gaussian kernel, FWHM 8 mm), trials were modeled at the first level as events of fixed duration (4 sec), and convolved with a standard BOLD hemodynamic function (as provided by the SPM software). To model the effect of preferences for individual items, we first computed the preference scores of each item as the number of times the item was chosen in the whole scanning session. We then computed in each trial the difference between the preference scores of the two items displayed in the trial, as an estimate of the gradient of preference in the trial, and their sum, as an estimate of the total appetitive salience of the trial. After centering, these computed values were entered as separate ‘parametric modulations’ of the trial regressor in the SPM software. Hence, the model included a regressor for the effect of trials relative to fixation, modeling stimulus encoding and response generation common to all trials, and the parametric modulations, modelling the interaction of this effect with the gradient of preference (for the score difference) and total appetitive salience (for the score sum). A second regressor included events for misses (if any). Realignment parameters were added to the model as confounding covariates. The fitted coefficients of the model (which included an AR(1) term for the residuals) were then brought to the second level to model individual variability as a random effect. To verify coverage of the ventromedial prefrontal cortex (which is prone to susceptibility artefacts), we checked the masks created by the SPM software based on sufficient tissue intensity values.

When participants respond to all trials and give consistent responses, these two regressors (difference and sum of the preference scores) are orthogonal (Viviani et al. 2018). This property may be difficult to prove in the general case, but it is easy to verify its validity for a specific number of items. In MATLAB,

> #possible range of scores for 6 items scores = 0:5; #possible arrangements of scores, taken two at a time trials = nchoosek(scores,2); #orthogonality of difference and sum of scores corrcoef([trials(:,2)-trials(:,1), trials(:,1)+trials(:,2)])

The possible pairings of scores occurs in a different order in each experiment, but the correlation is commutative with respect of the ordering of differences and sums of scores. The orthogonality makes precise the sense in which the effects of the differences of scores may be claimed not to be attributable to any property of the overall display, including its salience. The orthogonality does not obtain when participants miss trials, but since there were no participants with more than two misses, this effect is minor.

The signal course of Figure 3 was obtained by fitting a Fourier series of 6 basis functions for 7 time-points in the interval of the plot (fda software, MATLAB version, Ramsay and Silverman 2005) to the data residualized from the fit for the intercept, the regressor for the misses (if present), and the movement covariates. The trials were defined by an interval of 15 seconds after each onset, allowing selection of relevant scans and definition of time points of data collection (for details on this procedure, see Viviani et al. 2005). Ninety-five percent confidence intervals for the fitted curves were computed point-wise, as customary in functional data analysis (Ramsay and Silverman 2005).

Regions of interest were defined by the Jülich atlas for the amygdala (Amunts et al. 2005, obtained from the SPM anatomy toolbox, http://www.fz-juelich.de) and a bounding box in the orbitofrontal cortex at the coordinates reported in Levy and Glimcher (2012). Corrected significance levels were computed with permutation testing (Holmes et al. 1996, 8000 permutations). Cluster-level corrections for the whole brain were obtained from clusters defined by the a priori threshold *p* < 0.005 computed with a permutation method. Within regions of interest, the cluster definition threshold was set a priori to *p* < 0.01 to account for the small regional size. Overlay images were produced with the freely available software tool MriCroN (Chris Rorden, http://www.mccauslandcenter.sc.edu/mricro).

## Acknowledgments

This study was funded by a Neuron-ERANET grant (project BrainCYP, Grant number BMBF 01EW1402B) and by collaborative grants with the Federal Institute for Drugs and Medical Devices (BfArM, Bonn, Grants No. V-15981/68502/2014-2017 and V-17568/68502/2017-2020) to Roberto Viviani. The authors declare no conflict of interest.

